# Crossroads of assembling a moss genome: navigating contaminants and horizontal gene transfer in the moss *Physcomitrellopsis africana*

**DOI:** 10.1101/2023.10.30.564737

**Authors:** Vidya S. Vuruputoor, Andrew Starovoitov, Yuqing Cai, Yang Liu, Nasim Rahmatpour, Terry A. Hedderson, Nicholas Wilding, Jill L. Wegrzyn, Bernard Goffinet

## Abstract

The first chromosome-scale reference genome of the rare narrow-endemic African moss *Physcomitrellopsis africana* is presented here. Assembled from 73x nanopore long reads and 163x BGI-seq short reads, the 414 Mb reference comprises 26 chromosomes and 22,925 protein-coding genes (BUSCO: C:94.8%[D:13.9%]). This genome holds two genes that withstood rigorous filtration of microbial contaminants, have no homolog in other land plants and are thus interpreted as resulting from two unique horizontal gene transfers from microbes. Further, *Physcomitrellopsis africana* shares 176 of the 273 published HGT candidates identified in *Physcomitrium patens*, but lacks 98 of these, highlighting that perhaps as many as 91 genes were acquired in *P. patens* in the last 40 million years following its divergence from its common ancestor with *P. africana*. These observations suggest rather continuous gene gains via HGT followed by potential losses, during the diversification of the Funariaceae. Our findings showcase both dynamic flux in plant HGTs over evolutionarily “short” timescales, alongside enduring impacts of successful integrations, like those still functionally maintained in extant *Physcomitrellopsis africana*. Furthermore, this study describes the informatic processes employed to distinguish contaminants from candidate HGT events.

**Article Summary:** The first draft genome of the rare South African endemic moss *Physcomitrellopsis Africana* is presented. The 414 Mb assembly contains 22,925 genes, including two uniquely horizontally transferred genes, but lacks 97 of the microbial genes previously identified in the closely related model, *Physcomitrium patens* - highlighting the dynamic role of HGT in the evolution of these moss genomes and loss. This study presents best practices for contamination detection and new insights into HGT identification.

## Introduction

Horizontal gene transfer (HGT), also referred to as lateral gene transfer (Zhaxybayeva and Doolittle 2011) is the lateral movement of genetic material between distant branches of the tree of life. This process is ubiquitous among bacteria, facilitating rapid adaptation through exchange of ecologically important genes (Aminov 2011). While HGT is less common in eukaryotes than in prokaryotes, it plays a role in shaping eukaryotic evolution with about 0.04-6.49% of eukaryotic genes originating from HGT from microbes (Van Etten and Bhattacharya 2020). During the evolution of land plants, increasing interactions with rhizosphere microbes, particularly bacteria and mycorrhizal fungi, have enabled occasional horizontal transfer of functional genes between these distantly related lineages (Martin *et al*. 2017). For example, the gene for Killer Protein 4 (KP4) was likely acquired by mosses through HGT from ascomycete fungi (Guan *et al*. 2023). Similarly, the HET domain, common in fungal heterokaryon incompatibility genes and involved in self/non-self-recognition, was identified as a truncated form in *Physcomitrium patens* and may play a role in defenses against microbial pathogens (Sun *et al*. 2020). Van Etten and Bhattacharya (2020) suggested that rather than being merely anecdotal, HGT has been an important evolutionary force driving the adaptations of land plants to new habitats and stressors throughout their evolution, as previously already hypothesized by Yue *et al*. (2012).

Accurately detecting the taxonomic origin of genes and confirming horizontal gene transfer (HGT) events is challenging. Many reported cases of HGT in published genomes have been proven to be artifacts resulting from contamination that had gone undetected. For example, the initial report that 17% of the genes in the genome of the tardigrade *Hypsibius dujardini* originated from HGT (Boothby *et al*. 2015), has subsequently been revised to only 0.2% (Koutsovoulos *et al*. 2016). Similarly, claims of HGT in mammals also turned out to be erroneous due to bacterial contamination (Douvlataniotis *et al*. 2020), highlighting the need for careful analyses and experimental validation. Systematic screening of 43 arthropod genomes by Francois *et al*. (2020) revealed extensive bacterial contaminants, often outnumbering true horizontal gene transfer events. For example, in the bumblebee *Bombus impatiens*, most contaminating genes were concentrated on just 30 contaminant scaffolds. Based on the size and number of contaminating sequences, it was concluded that the genome of the symbiont *Candidatus Schmidhempelia bombi* was co-assembled with that of its host. Strategies to confidently identify HGT, including taxonomic assignment of candidates via BLASTp/DIAMOND, phylogenetic analysis with candidate and donor proteins, synteny analysis of flanking genes, and quantitative PCR validation, have been presented to address these issues (Francois *et al*. 2020).

*Physcomitrellopsis africana* inhabits transitional zones between grassland and forests in coastal habitats in South Africa. It is currently the sole species of the genus, which likely should also include several species of the paraphyletic genus *Entosthodon* (Wilding 2015). Its genome of 414 MB, the first genome of an African bryophyte, is presented here. This resource complements those available for *Funaria hygrometrica* and *Physcomitrium patens* (Rensing *et al*. 2008; Kirbis *et al*. 2022) and thereby constitutes a fundamental resource to further the reconstruction of the evolution of the genome of the model, *Physcomitrium patens* (Rensing *et al*. 2020). Through rigorous contamination screening and verification, two genes emerge as candidates resulting from unique HGT events. Informatically validating HGT events in *Physcomitrellopsis africana* yields evolutionary insights while underscoring the need for stringent standards to support HGT conclusions from genomics data.

## Materials and Methods

### Sample collection and culturing

A population of *Physcomitrellopsis africana* was sampled in October 2010 from a coastal forest in the Eastern Cape Province of South Africa (Dwesa National Park, along trail to chalets behind campground, coordinates 32 °18.222’ S 28 °49.666’, at ± sea level). The voucher specimens collected by Goffinet (collection numbers 10326 and 10329, with J. Goffinet, T Hedderson and N Wilding) are deposited in the CONN herbarium under accession numbers CONN00235389 and CONN00235388, respectively. Specimen 10326 (culture and long read library DNA #5074) provided DNA for long read sequencing, whereas 10329 (RNA and BGI-Seq DNA #5075) provided RNA and DNA for short read sequencing. Sterile cultures were first established on Knop medium using spores from a single operculate capsule. The gametophytes were harvested, ground, and spread on a rich sandy loam soil in PlantCon tissue culture containers (MP Biomedicals, Solon, OH, USA), and maintained in a growth chamber under 16 h of daylight at about 24 °C.

### Genomic DNA and RNA extraction

Gametophytic tissue consisting of stems and leaves, of *Physcomitrellopsis africana* was harvested from fresh soil cultures under a dissecting microscope and ground in liquid nitrogen. DNA was extracted by following a modified protocol by Young (2022). The quality of the DNA sample 5074 was assessed by quantitative PCR prior to sequencing, yielding a DIN score of 7.0 and concentration of 40.9 ng/μL.

DNA for short read sequencing was extracted using the NucleoSpin Plant midi DNA extraction kit, following the manufacturer’s protocol (Macherey-Nagel, Düren, Germany). DNA quality was evaluated using a Qubit® 3.0 Fluorometer (Thermo Fisher Scientific, USA). Total RNA was extracted from approximately 1 g of fresh gametophytic tissue using the RNeasy Plant Mini Kit (Qiagen, Valencia, CA, USA).

### Genome and transcriptome library preparation and sequencing

DNA was prepared for long-read PromethION sequencing through a DNA repair step, generating blunt ends, and ligating sequencing adapters followed by priming of the flow cell, as described in the Oxford Nanopore Technologies Amplicons by Ligation (SQK-LSK109) protocol. The HMW DNA library was sequenced on an Oxford Nanopore PromethION (Center for Genome Innovation, UConn) using a FLO-PRO0002 flow cell.. The two short-read DNA libraries were prepared following the methods used by Yu et al (2020), and was sequenced (150 bp PE) on two lanes of a BGISEQ-500 (BGI-Shenzhen, China).

Approximately 1 μg of RNA was used to generate a paired-end library with an insert fragment size of 200–300 bp of the corresponding cDNA. RNA purification, reverse transcription, library construction and sequencing were performed at WuXi NextCode (Shanghai, China). The captured coding regions of the transcriptome from total RNA were prepared using the TruSeq® RNA Exome Library Preparation Kit. The two RNA libraries were sequenced on one lane of an Illumina HiSeq 2000 (100 bp PE) at the WuXi NextCode (Shanghai, China).

### Quality control of genomic and transcriptomic reads

The genomic short reads were first assessed with FASTQC v.0.11.7 (Andrews 2010). In preparation for assembly with Haslr and Wengan, Sickle v.1.33 with a minimum quality score threshold of 30 (-q) and a minimum length of 50 bp (-l) was employed for read trimming. Nanoplot v.1.21.0 was used to quantify and assess the quality of the nanopore reads. To detect potential contamination, the long reads were aligned against the pre-indexed bacterial, human and viral databases with a metagenomic classifier, Centrifuge v.1.0.4-beta (p+h+v; min-hit length was increased to 50 bp) (Kim *et al*. 2016). Reads aligning to the database were removed. The quality of the transcriptomic short reads was assessed with FASTQC v.0.11.7 (Andrews 2010). The reads were trimmed with Sickle v1.33 (Joshi and Fass 2011) with a minimum quality score threshold of 30 (-q) and a minimum length of 40 bp (-l).

### Genome size estimation

A single lane of short-read genomic data from accession 5075 was employed to estimate the genome size. The k-mer distribution was calculated using Jellyfish v2.2.6 (Marçais and Kingsford 2011) and size estimates were processed with GenomeScope v2.0 (Ranallo-Benavidez *et al*. 2020; Supplementary Fig. S1).

### Transcriptome assembly

The transcriptome was independently assembled to provide protein-level evidence for the structural annotation of the genome, using Trinity v2.6.6 (Grabherr *et al*. 2011), with a minimum contig length of 300bp. Contigs with minimal read support, post assembly, were removed (FPKM > 0.5) with RSEM v1.3.0 (Li and Dewey 2011). Transdecoder v3.0.1 (Haas and Papanicolaou 2011) was used to translate the remaining contigs into Open Reading Frames (ORFs) and remove sequences without a viable frame. To aid Transdecoder in the identification of ORFs, searches against the Pfam database were performed with HMMER v3.1b2 (Zhang and Wood 2003). Transdecoder annotated the putative transcripts as complete, partial, and internal. Those without a defined start and stop codon (defined as internals) were removed (split_frames.py). The final set of peptide sequences were functionally annotated with EnTAP v0.8.0 (Hart *et al*. 2020) against NCBI’s nr protein database and the UniProt/Swiss-prot reference databases. EnTAP was run with contaminant filters that included bacteria, archaea, fungi, and insecta. Transcripts with high confidence alignments to these organisms were removed (contam_removal.py).

### Genome assembly

Hybrid genome assembly, integrating both long and short-read data, was conducted with MaSuRCA v.4.0.3 (Zimin *et al*. 2013), Wengan v.0.2 (Di Genova *et al*. 2021), and Haslr v.0.8a1 (Haghshenas *et al*. 2020). Additionally, a separate assembly using only long-reads as input was conducted with Flye v.2.5 (with three polishing iterations) (Kolmogorov *et al*. 2019).

The Flye, Wengan, and Haslr assemblies were polished following long-read alignment with Medaka v1.3.2 (github.com/nanoporetech/medaka). Post assembly with Wengan, the assembly was filtered to remove scaffolds less than 3 Kb. To further improve the accuracy of the assemblies, the hybrid assembly generated by MaSuRCA was polished with the short reads using Pilon v.1.24 (Walker *et al*. 2014). Subsequently, the selected MaSuRCA assembly was processed with Purge Haplotigs v.1.0 (Roach *et al*. 2018).

The Purge Haplotigs pipeline categorized the assembled scaffolds into four coverage levels based on the distribution of mapped reads. This categorization enabled the identification and removal of redundant sequences exhibiting low coverage, presumed to represent erroneous duplicates given the haploid genome. Specific cutoff values of 0, 7, and 65 for the coverage levels were selected to delineate scaffolds to be retained or discarded, based on read depths (Supplementary Fig. S1). The term “allele” is used to describe these redundant sequences for convenience, although technically inaccurate for a haploid genome. The coverage analysis and purging allowed isolation of the primary genome sequence from duplications and artifacts generated during assembly.

To evaluate the quality of the assemblies, QUAST v5.2.0 (Gurevich *et al*. 2013) and BUSCO v4.1.2 (viridiplantae_odb10:425 single copy orthologs) (Manni *et al*. 2021) were employed. Each assembly was also evaluated with Merqury v1.3 (Rhie *et al*. 2020).

### Genome annotation

#### Repeat library construction and masking

The repeat library for the final MaSuRCA assembly was generated using RepeatModeler v.2.01 (Flynn *et al*. 2020) with the long terminal repeat (LTR) discovery pipeline enabled. The genome was then soft-masked with RepeatMasker v.4.0.9-p2 using the consensus repeat library from RepeatModeler (Smit *et al*. 2013-2015).

#### Structural and functional genome annotation

The RNA reads were aligned to the soft-masked MaSuRCA assembly with HISAT2 v.2.1.0 (Kim *et al*. 2019) to provide evidence for protein-coding gene prediction. Two gene prediction analyses were run on the soft-masked assembly using BRAKER v.2.1.5 (Brůna *et al*. 2021), one with RNA-Seq alignment evidence and one with protein evidence originating from *de novo* assembled transcriptome. Gene predictions from both BRAKER runs were integrated with TSEBRA v.1.0.3 (Gabriel *et al*. 2021). From this point, separate assessments were conducted on the RNA-Seq evidence gene predictions (BRAKER) and the final TSEBRA gene predictions to select the best approach. Putative genes were removed from both sets if they did not contain a complete protein domain. This filter was applied with Interproscan v.5.35-74.0 (Jones *et al*. 2014) using the Pfam database v32.0 (Finn *et al*. 2014). It is worth noting that mono-exonic genes can be the result of fragmented annotations and the target metric of 0.2 (mono:multi-exon gene ratio) is often achieved through protein domain filters (Vuruputoor *et al*. 2022). Metrics for the gene predictions were generated with AGAT (Dainat 2020) and BUSCO. After assessment, the filtered BRAKER gene predictions were selected for functional annotation with EnTAP v.0.10.8 (Hart *et al*. 2020). Functional annotation reports from EnTAP (both sequence similarity search and EggNog taxonomy scope classifications) allowed for the identification of non-target species scaffolds in the assembly (Huerta-Cepas *et al*. 2019).

#### Assembly-level contaminant filtering

Using the functional annotation results from EnTAP, contaminated scaffolds were removed. Scaffolds with a length of 10 Kb or less, and with 40% or more of their total genes classified as archaea, bacteria, or fungi, were removed. Additionally, scaffolds greater than or equal to 10 Kb with 55% or more genes classified as archaea, bacteria, or fungi, were also excluded. The final annotation was then assessed for the annotation rate using EnTAP, the mono:multi ratio using AGAT, and BUSCO completeness.

#### Horizontal gene transfer candidate identification

To identify candidate HGTs in *Physcomitrellopsis africana*, protein sequence similarity searches were conducted with Diamond v2.1.8 (Buchfink *et al*. 2021). The protein sequences of *Physcomitrellopsis africana* were aligned against “donor” databases, which included sequences from bacteria, fungi, archaea, and metazoa from NCBI’s nr database. Additionally, the same proteins were aligned to “recipient” databases containing sequences from Streptophyta, Tracheophyta, Embryophyta, Viridiplantae, and Spermatophyta (Ma *et al*. 2022). Although these categories are not fully exclusive, each database was utilized separately to systematically assess presence across plants at different evolutionary divergence points.

To identify candidate genes representing putative horizontal gene transfer (HGT) events unique to *Physcomitrellopsis africana*, the following criteria were utilized: genes were required to have between one and four significant sequence alignments (E-value <1e-5) to microbial donor databases, while exhibiting no significant sequence similarity to plant recipient databases. This range of one to four microbial alignments was selected to capture potential HGTs while avoiding ubiquitous domains shared across many microbes. The lack of hits to plant databases was intended to enrich for *Physcomitrellopsis africana*-specific sequences, rather than those conserved across plants through vertical inheritance. At this stage, any scaffolds containing only bacterial or fungal genes, without any plant-related genes, were removed from the assembly.

To assess the validity of the two proposed HGT candidates, the target proteins were independently aligned to the reference genome with Miniprot v0.7. To examine independent transcriptomic support, the RNA reads were aligned to the genome with HISAT2 v2.1.0 and assembled via StringTie2 v2.2.1 (Kovaka *et al*. 2019).

#### Analyzing HGT candidates from Physcomitrium patens

The 273 putative horizontally transferred genes (HGTs) previously identified in *Physcomitrium patens* (Yue *et al*. 2012; Ma *et al*. 2022) were independently searched against the *Physcomitrellopsis africana* and *F. hygrometrica* protein sets using DIAMOND v2.1.8 (Buchfink *et al*. 2021). DIAMOND searches were conducted with an E-value cutoff of 1e-5 and max target sequences set to 1. Hits against *Physcomitrellopsis africana* and *F. hygrometrica* were collected and merged to generate a summary table with *Physcomitrium patens* HGTs and the respective top hits in each species.

### Scaffolding the genome to chromosome scale

A single HiC library was prepared for *Physcomitrellopsis africana* using Proximo Hi-C kit (Phase Genomics) and sequenced as paired-end reads on an Illumina NovaSeq platform (Center for Genome Innovation, UConn). The reads were aligned to the MaSuRCA genome assembly using BWA v0.7.17 but did not provide sufficient coverage due to contamination. Instead, the *Physcomitrium patens* T2T reference genome assembly (Bi *et al*. 2024) was used to scaffold the *Physcomitrellopsis africana* assembly to chromosome-scale with RagTag v2.1.0 under default parameters, except a minimum fragment size threshold of 1 Kb was specified.

### Comparative genome analyses

A comparative analysis of the protein-coding gene space was conducted with OrthoFinder v.2.5.1 (Emms and Kelly 2019) with *F. hygrometrica* (Kirbis *et al*. 2022) and *Physcomitrium patens* v3 (Lang *et al*. 2018). To provide a preliminary estimate of gene family size dynamics, gene counts from each species in the assembled orthogroups were categorized as neutral, expanded, or contracted. The first and third quartiles were calculated for the distinct gene counts within a gene family for each species. If the number of genes from a species was lower than the first quartile or higher than the third quartile, then the gene family was categorized as “contracted” or “expanded”, respectively. If the number of genes did not fit with either of these two criteria, then the gene family was considered “neutral”. The longest gene for each orthogroup was used to assign functional attributes to all genes in the group from the original EnTAP annotation. If the longest gene did not originate from *Physcomitrellopsis africana*, then the functional annotation was derived from either *Physcomitrium patens* or *F. hygrometrica*.

GOSeq (Young *et al*. 2010) enrichment analysis was performed in R v4.2.0. GO terms were extracted for each gene from the EnTAP run. Enrichment analysis was investigated separately for the *Biological Process* and *Molecular Function* GO categories. Paralogs of Light Harvesting Complex (LHC), Serine/Threonine-protein kinase (STN) 7, and STN8 were identified with Diamond v2.8.1 (E-value <1e-5).

### Whole genome duplication analysis

Chromosome-scale genomes of *F. hygrometrica* and *Physcomitrium patens* were assessed with the reference genome generated for Ph*yscomitrellopsis africana*, with wgd (v.2.0.19) to characterize whole genome duplication events (Chen and Zwaenepoel 2023). Each species was compared against itself. Nucleotide sequences (CDS) were used as input to Blast & Markov clustering (MCL) (Altschul *et al*. 1997; van Dongen 2000). A Ks distribution was constructed using the ‘ksd’ subcommand and ‘mcl’ output (Yang 2007; Katoh and Toh 2008; Price *et al*. 2010). Next, a collinearity analysis was performed with ‘syn’ using the structural genome annotations (Proost *et al*. 2011). Gaussian mixture models, via ‘mix’, were used to generate Ks distributions to aid in component interpretation. Model fit was evaluated with the Bayesian and Akaike information criterion (BIC/AIC).

## Results and Discussion

### Sequencing and quality control of genomic reads

The single ONT run generated 16 million reads (N50: 7,518 bp; 127X coverage: 404 Mb; 116X coverage: 440 Mb; Supplementary Table S1). Centrifuge filtering reduced this set to approximately 14M reads (N50: 4,561 bp; 80X coverage: 404 Mb; 73X coverage: 440 Mb). The primary contaminants included bacteria from *Xanthomonadaceae* (10.68%) followed by *Bradyrhizobiaceae* (0.7%). The short-read genomic libraries (100 bp PE) generated 529 M reads. Following trimming with Sickle, 360 M reads remained (178X coverage: 404 Mb; 163X coverage: 440 Mb; Supplementary Table S2). Coverage estimates are provided for both the original estimate (404 Mb) and the final assembled genome size (440 Mb).

### Transcriptome assembly

A total of 36.58M RNA short reads were generated. Following quality filtering using Trimmomatic, the dataset was reduced to 31.84M reads. The *de novo* assembly process with Trinity yielded 143,150 contigs. After expression filtering via RSEM, the number of contigs was reduced to 123,373. Identifying open reading frames (ORFs) in the contigs resulted in 110,852 successfully translated transcripts. The average sequence length of these translated ORFs was 803 bp, with an N50 value of 1,038 bp. To enhance the quality of the transcriptome assembly, internal sequences and putative contaminants were removed, resulting in 74,997 total transcripts. After removing internal sequences and contamination, the total unique sequences with a sequence similarity search alignment are 51,982 (69.3%). An assessment of contamination revealed that 8,720 transcripts, 16.78% of the transcriptome, originated from various sources such as amoeba (0.06%), bacteria (0.64%), fungi (15.93%), or insecta (0.12%), as identified by EnTAP. A further 15,500 (18.5%) sequences remained unannotated. The final BUSCO score of the remaining transcripts was C:82%[D:35.7%]. A total of 30,657 (36.6%) transcripts aligned to *Physcomitrium patens* proteins.

### Genome assembly

#### Initial assembly

Multiple genome assembly approaches were employed to generate comprehensive draft assemblies of the *Physcomitrellopsis africana* genome (Fig. 2). The long-read approach, Flye, assembled 538.68 Mb in 8,388 scaffolds, with an N50 of 152.58 Kb. The BUSCO completeness score was C:84.0%[D:12.2%] and the Merqury QV score was 21.1. Among the hybrid approaches, Haslr produced a 295.98 Mb reference distributed across 12,738 scaffolds, with an N50 of 52.29 Kb. The BUSCO score was C:94.1%[D:13.2%], and the QV score was 11.9. Wengan assembled a total of 466.64 Mb across 10,516 scaffolds, with an N50 of 119.42 Kb a BUSCO score of C:96.4%[D:18.8%], and a QV score of 27.4. Finally, MaSuRCA assembled 506.22 Mb across 3,571 scaffolds, with an N50 of 381.33 Kb, (BUSCO: C:96.4% [15.5%], and a QV score of 31.0).

#### Polishing and improving genome assembly

The Medaka polished Flye assembly had a genome size of 530.57 Mb in 8,386 scaffolds, and the N50 was decreased to 145.17 Kb. The BUSCO dropped to C: 90:3%[D:11.5%]. The QV score decreased to 19.6. Polishing of the Haslr assembly resulted in a genome size of 295.73 Mb, across 12,737 scaffolds. The N50 remained almost unchanged at 52.31 Kb, and the BUSCO score reduced to C:90.6%[D:9.4%]. The QV score more than doubled to 25.2. Filtering for scaffolds less than 3 Kb, followed by Medaka, resulted in a smaller genome size for the Wengan assembly at 434.41 Mb across 7,639 scaffolds. The N50 increased to 132.40 Kb. The BUSCO score was reduced to C:93.2%[D:11.8%], and the QV increased slightly to 27.5.

Polishing with Pilon had very minimal influence on the completeness, as expected, and substantial impact on the accuracy of the MaSuRCA assembly. The polished MaSuRCA hybrid assembly size increased slightly to 507.10 Mb across 3,590 scaffolds, with a slightly decreased N50 of 378.43 Kb. The BUSCO completeness remained the same at C:96.4%[D:15.5%], and the QV score increased to 35.6.

#### Refinement of genome assemblies with Purge-haplotigs

The MaSuRCA assembly was selected for further refinement. This decision was based on the overall quality assessed by Merqury, BUSCO completeness, and overall contiguity. The MaSuRCA assembly was refined with Purge-haplotigs, which reduced the assembly length to 502.34 Mb across 3,237 scaffolds with an N50 value of 382.25 Kb. The BUSCO completeness score remained the same (e.g., 96.4%) and the QV score was minimally reduced to 35.5. At 502.34 Mb, the assembled genome is ∼100 Mb longer than the k-mer based estimate (440 Mb, Supplementary Fig. S2).

### Genome annotation

#### Repeat identification

RepeatModeler produced a library containing 580 unique repeats that were used to softmask 50.22% of the final assembly (Supplementary Table S3). Of the repeats, 36.53% were composed of *Ty3/Gypsy* and 1.46% of *Ty1/Copia*. This is similar to the pattern in *Physcomitrium patens*, wherein approximately 57% of the genome is composed of repeat elements, with long terminal repeats (LTRs), particularly the *Gypsy* family, accounting for 48% of the masked genome (Lang *et al*. 2018). These findings align with observations by Kirbis *et al*. (2022), suggesting a common pattern regarding the activation of *Gypsy* elements in *Physcomitrellopsis africana* and *Physcomitrium patens*. In contrast, *F. hygrometrica*, which diverged from *Physcomitrium patens* 60 to 80 MYA (Medina *et al*. 2018; Bechteler *et al*. 2023), exhibits a lower overall repeat estimate of 35%. Here, *Gypsy* elements contribute less to the LTR content, i.e., roughly 10%, whereas *Copia* elements contribute 17%.

#### Protein-coding gene identification

The *Physcomitrellopsis africana* RNA-Seq library had an overall alignment rate of 78.43%, likely due to contaminant content in the source extraction. Regardless, a substantial set of 37M reads were retained, exceeding the minimum needed for prediction. The first BRAKER2 predictions, using RNA-Seq evidence alone, generated 60,917 protein-coding genes with a BUSCO completeness score of C:95.8%[D:15.3]. The mono:multi exonic ratio was 0.52. With only protein evidence, BRAKER2 generated 37,752 gene predictions with a BUSCO score of C:46.1%[D:16.0]. Merging these predictions, as recommended by TSEBRA, resulted in a set of 45,737 genes with a BUSCO score of C:91.3%[D:15.1], and a mono:multi exonic ratio of 1.06. The gene prediction set generated with RNA-Seq alignment evidence exclusively was selected for further refinement due to its higher BUSCO score and lower mono:multi ratio compared to the merged TSEBRA transcripts. Through protein domain filtering (InterProScan filter), the number of mono-exonic genes was further reduced from 20,081 to 12,696, producing a total of 23,561 genes (BUSCO: C:88.5%[D:13.8%]; mono:multi ratio: 0.08).

Based on the functional annotations of the gene space, we removed 831 scaffolds and 1250 genes related to contaminants. This also reduced the assembly length to 440 Mb in 2,406 scaffolds with an N50 of 363 Kb, and an assembly BUSCO score of C:96%[D:13.9%]. This substantial reduction led to an assembly within ∼40 Mb of the original k-mer-based estimate (e.g., 404 Mb).

The annotated protein-coding gene space in the contaminant-filtered assembly included 1,708 mono-exonic and 21,853 multi-exonic genes. A BUSCO score of C:93.8%[D:13.6%] and mono:multi ratio of 0.08 was reported. Reciprocal BLAST conducted by EnTAP produced an annotation rate of 78%, of which 93.6% aligned to *Physcomitrium patens* (Table 1; File S1). This contrasts with the annotation rate of 50% in *F. hygrometrica* (Kirbis *et al*. 2022), attributed to significant divergence time from *Physcomitrium patens,* and the lack of closely related species in public genomic databases (Kirbis *et al*. 2020; Rahmatpour *et al*. 2021). The higher annotation rate in this study suggests that many genes found in *Physcomitrium patens,* but not *F. hygrometrica,* may have been acquired in the ancestor of the *Physcomitrellopsis*-*Entosthodon*-*Physcomitrium* clade (Medina *et al*. 2019).

**Table 1.**
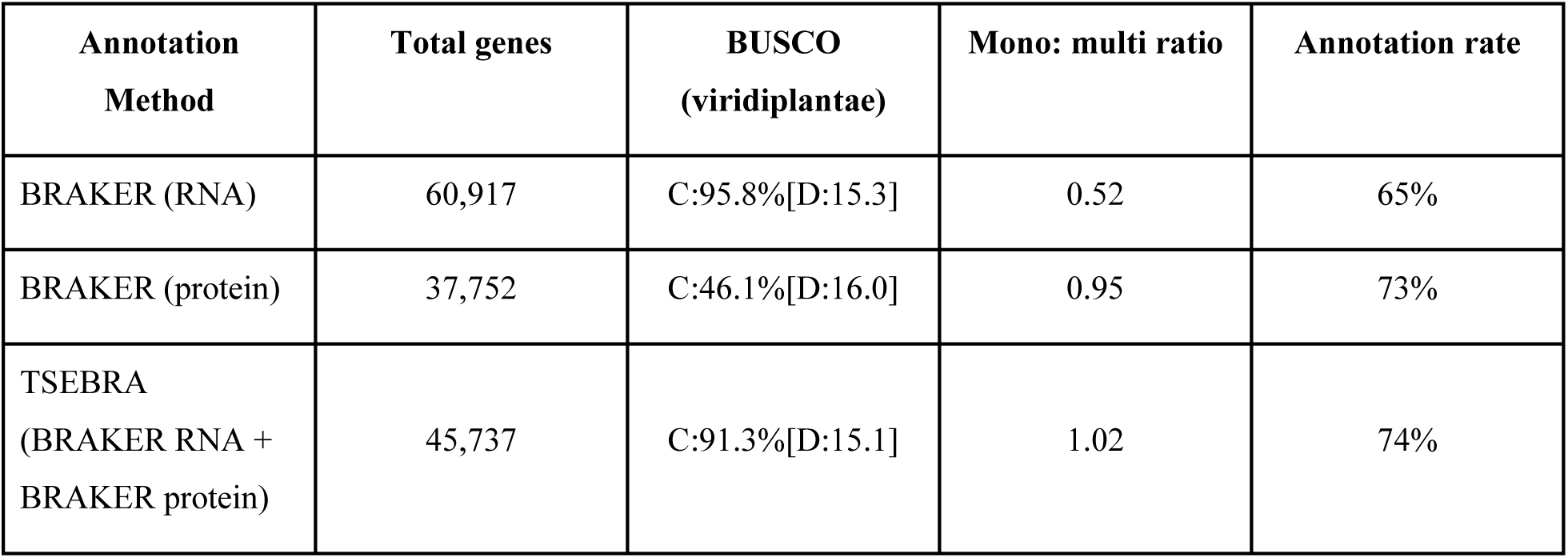
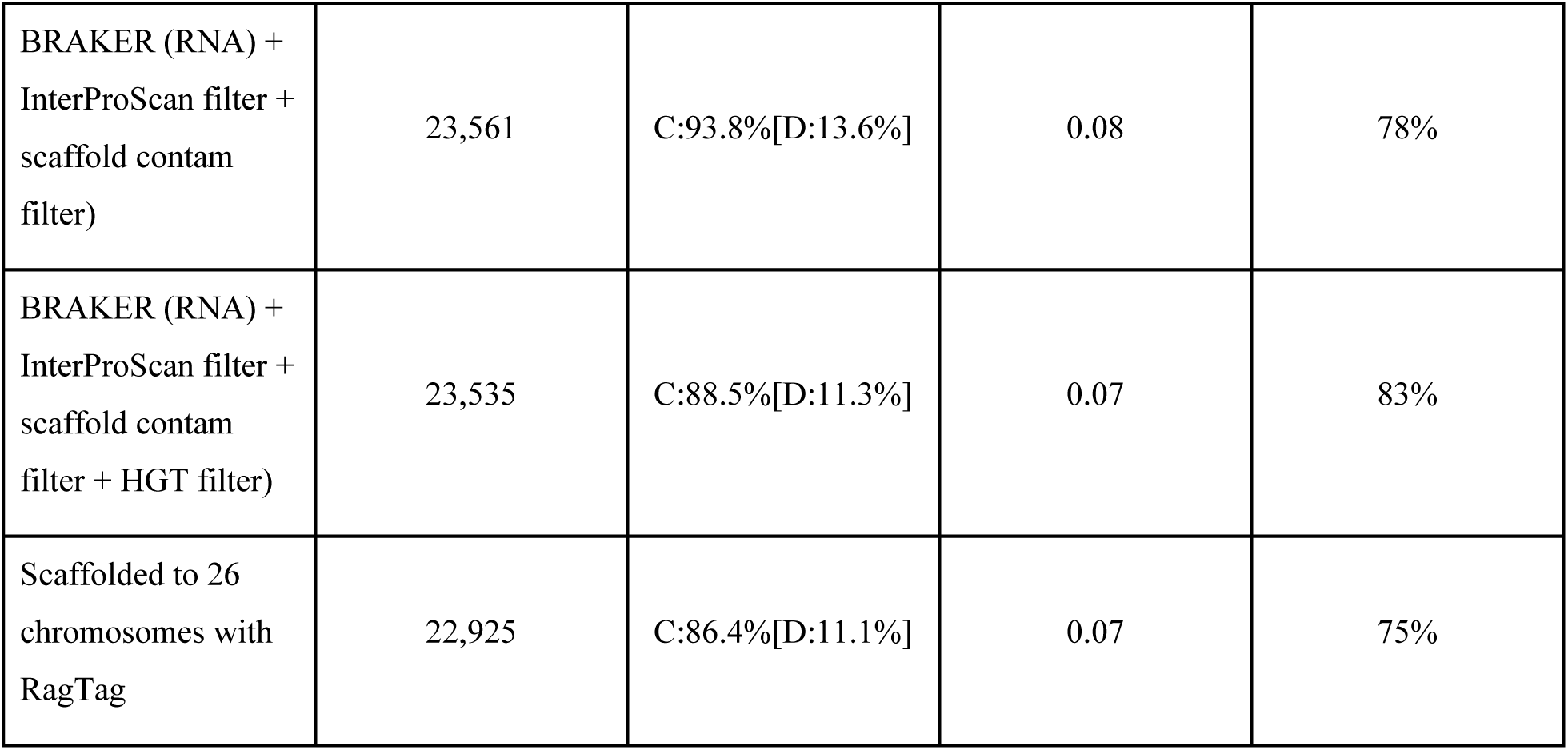
Genome annotation statistics for *Physcomitrellopsis africana*.

### Scaffolding the genome assembly to chromosome-scale

The T2T assembly of *Physcomitrium patens* (V4; Bi *et al*. 2024) was used to scaffold the *Physcomitrellopsis africana* genome assembly. The final assembly contained 440.65 Mb in 583 scaffolds with an N50 length of 15.04 Mb. The 26 largest pseudomolecules, corresponding to the *Physcomitrium patens* V4 haploid chromosome number, covered 414.21 Mb or 94% of the 440 Mb assembly. From here on, we will refer to these pseudomolecules as chromosomes, numbered according to their lengths. The final genome assembly had a BUSCO completeness of C:94.8% [S:80.9%, D:13.9%], F:1.2%, M:4.0%, n:425, and a total number of 22,925 protein coding genes. While most of the annotated gene space was retained, the BUSCO duplication score remained at 13%, compared to 11–15% for the *F. hygrometrica* and *Physcomitrium patens* genomes (Kirbis *et al*. 2022). Of the 557 contigs/scaffolds that did not contribute to the 26 chromosomes, 185 (33.34%) contained at least one protein coding gene annotation from the first annotation. Among those, three (1.6%) contained at least one gene that was annotated as a contaminant but did not meet the original thresholds that removed full scaffolds.

### Comparative genome analysis

The comparison of the genomes of *Physcomitrellopsis africana*, *F. hygrometrica*, and *Physcomitrium patens*, reveals that they share 13.5K orthogroups, but also hold 187, 939, and 958 unique orthogroups, respectively. *F. hygrometrica* and *Physcomitrellopsis africana* exclusively share more (1053) orthogroups than *Physcomitrellopsis africana* and *Physcomitrium patens* (745) (Fig. 3A), despite the latter sharing a more recent unique common ancestor (Fig. 1). Thus, whereas Funariaceae share a rather conserved architecture of their vegetative body even after at least 60 MY of divergence, their gene space varies considerably (Kirbis *et al*. 2022). Such differentiation was previously noted based on highly diverging transcriptomes (Rahmatpour *et al*. 2019), and interpreted as reflective of strong ecophysiological adaptations, which may be the driving force in bryophyte evolution (Glime 1990). Signatures of two ancient whole genome duplication (WGD) events were found in both *Physcomitrium patens* and *F. hygrometrica*. Previous studies have shown that genes related to regulation were preferentially retained after the first WGD in *Physcomitrium patens* (Lang *et al*. 2018). Our analysis of *Physcomitrellopsis africana* also retrieved two peaks, in agreement with the two WGD events in *F. hygrometrica* and *Physcomitrium patens*. The Ks peaks were at 0.56 and 0.92, and the AIC and BIC values accorded with two WGD events. To assess the gene expansion and contraction analysis, of the putative gene families from the *Physcomitrellopsis africana* annotation, 809 and 1,694 were, respectively, categorized as expanded and contracted (File S2). Enrichment analysis examined through Gene Ontology’s *Biological Process* category revealed 167 expanded terms and six contracted terms. In contrast, within the *Molecular Function* GO category, 62 were contracted, and no terms were significantly expanded.

**Fig. 1.**
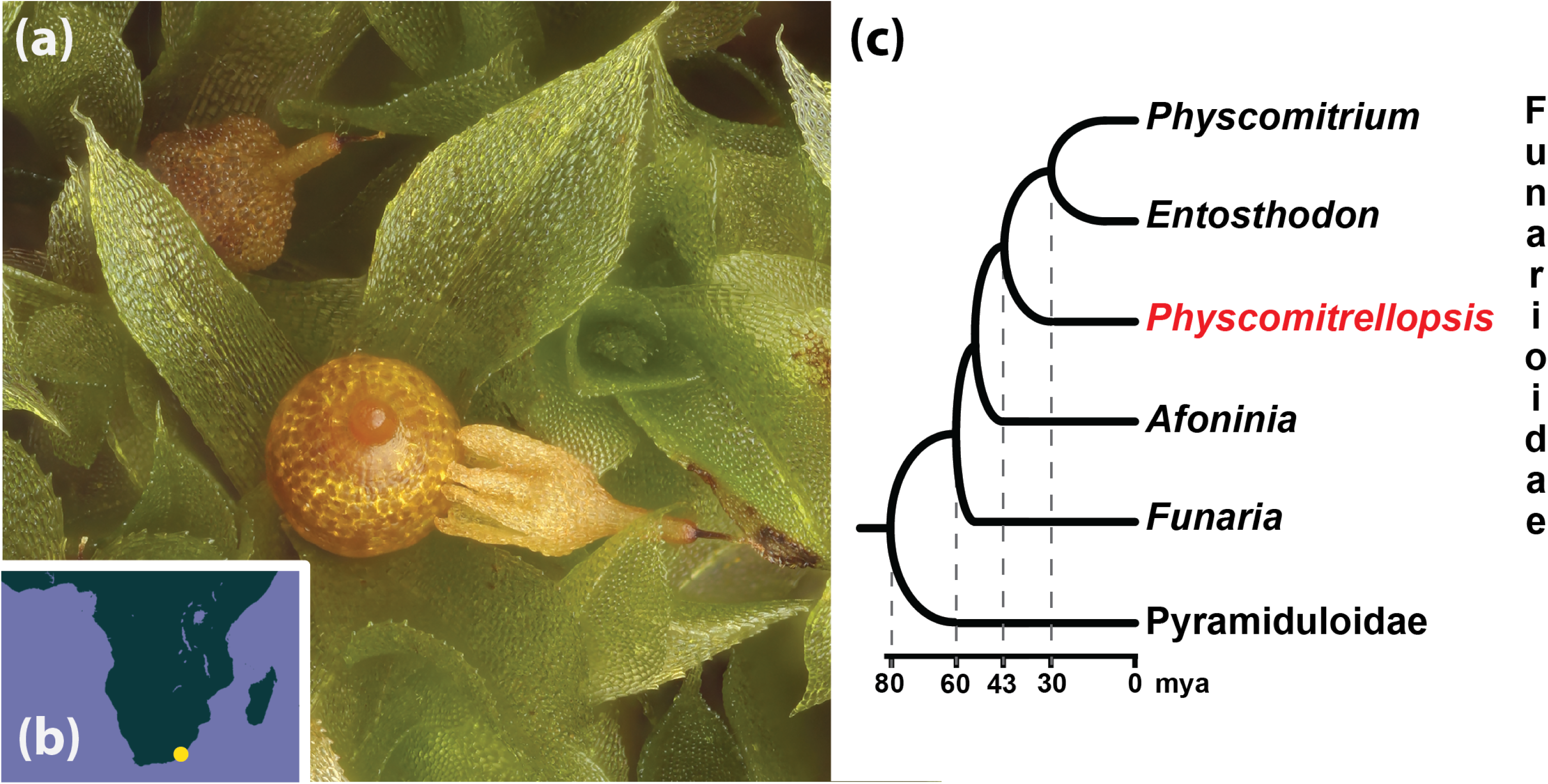
a) *Physcomitrellopsis africana* exhibits a reduced architectural complexity of the sporophyte similar to that observed in *Physcomitrium patens*, namely a sessile, aperistomate and cleistocarpous sporangial capsule. b) The known geographic distribution of *Physcomitrellopsis africana*, a rare narrow endemic to the Eastern Cape Region in South Africa. **C** Phylogenetic relationships and chronology of the evolution of *Physcomitrellopsis* based on Medina *et al*. (2018).

**Fig. 2.**
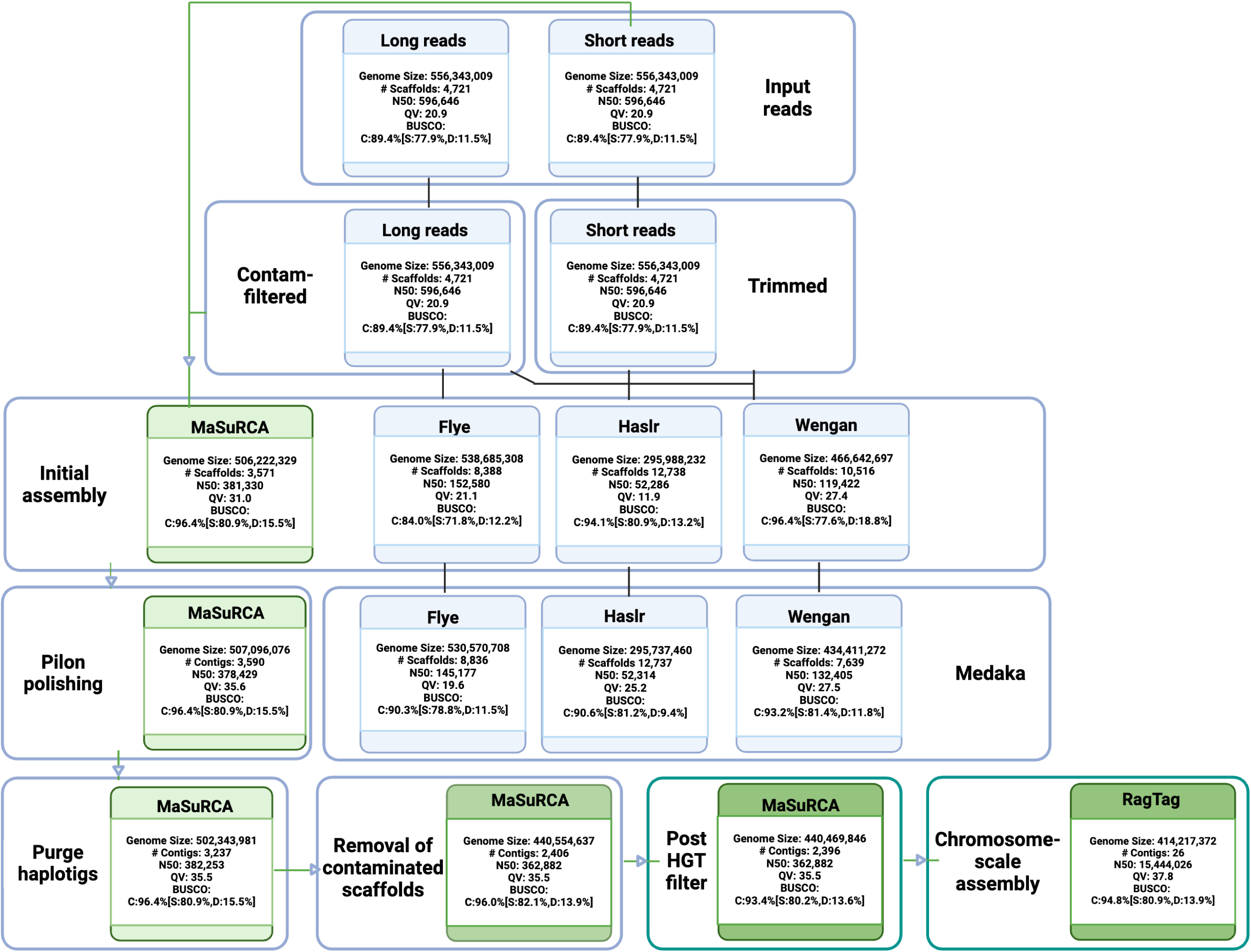
Workflow and statistics for read QC, genome assembly, polishing, and haplotype phasing. Short reads generated from BGI-seq were subject to trimming and the long reads (ONT) were filtered for contaminants with Centrifuge. Filtered and trimmed reads were utilized as input for four different genome assembly software tools: Flye (long-read only), Haslr, Wengan, and MaSuRCA. The green box indicates the selection of the MaSuRCA assembly for further analysis. The final assembly was polished using Pilon and phased with Purge haplotigs. A total of 831 scaffolds were removed as a result of contaminant filtering using EnTAP following structural genome annotation. Ten additional scaffolds were removed after HGT candidate assessment. The assembly was then scaffolded against the *Physcomitrium patens* V4 genome using RagTag, capturing 94% of the total assembly across 26 chromosomes. Summary statistics derived from Quast (N50 - scaffold/contig length at which 50% of the genome is contained in scaffolds/contigs of this size or greater), BUSCO, and Merqury (QV - Quality Value, an assessment of assembly contiguity) are displayed for each process.

**Fig. 3.**
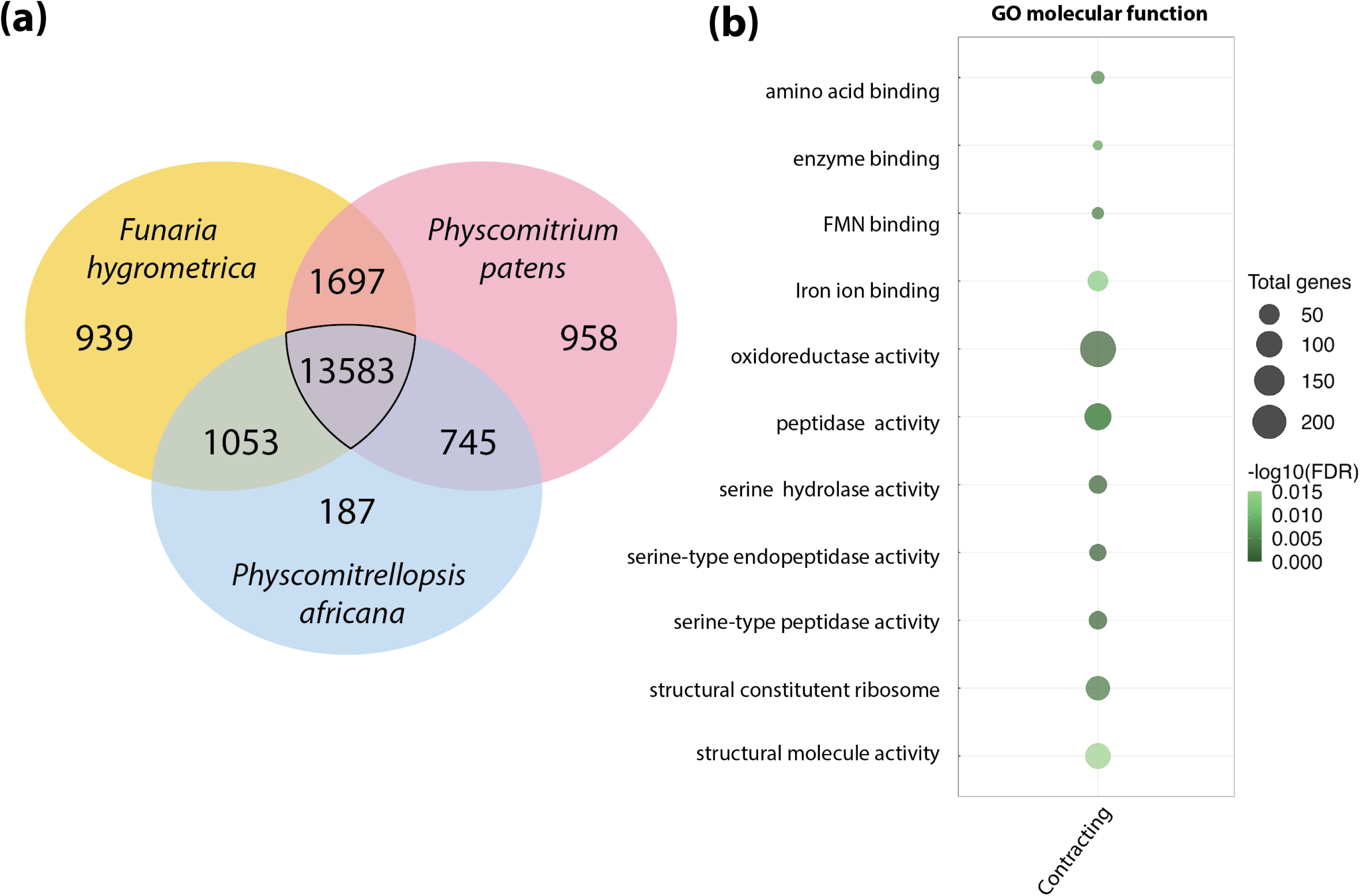
a) The total shared and unique orthogroups among *Funaria hygrometrica*, *Physcomitrellopsis africana*, and *Physcomitrium patens*. b) Enriched *Molecular Function* GO terms for gene families in a comparative analysis between *Physcomitrellopsis africana*, *Physcomitrium patens*, and *F. hygrometrica,* showed contraction in the GO terms related to oxidoreductase activity, as well as serine peptidase activity and FMN binding for *Physcomitrellopsis africana*. The size of each bubble represents the number of gene families within an orthogroup, and the gradient of the color denotes the significance level of enrichment-the darker green denotes more significance.

Although expanded *Biological Process* GO terms were excessively broad, orthogroup analysis revealed a pattern of contraction in GO terms pertaining to oxidoreductase activity, FMN binding, and serine protease activity (Fig. 3B). Whether this unique suite of downregulated categories diagnoses photosynthetic properties of *Physcomitrellopsis africana* only or of the expanded genus sensu Wilding (2015) remains to be tested. The observed changes in GO term enrichment related to oxidoreductase activity, FMN binding, and serine protease activity suggest potential differences in photosynthetic processes between *Physcomitrellopsis africana* and other moss species.

The complement of light-harvesting complex (LHC) genes is expanded in *Physcomitrium patens* compared to algae and vascular plants (Alboresi *et al*. 2008; Iwai *et al*. 2018). LHC proteins bind chlorophylls and carotenoids to facilitate light absorption and energy transfer to the reaction centers of Photosystems I and II. The LHC genes are classified into two groups: Lhca encodes antenna proteins for PSI (LHCI) whereas Lhcb encodes antenna proteins for PSII (LHCII). Although ancestral land plants contain several LHC homologs, further expansion occurred in *Physcomitrium patens* after whole genome duplication events (Alboresi *et al*. 2008; Rensing *et al*. 2008; Zimmer *et al*. 2013; Sun *et al*. 2023;). This led to a larger repertoire of LHC genes compared to the alga *Chlamydomonas reinhardtii* and the vascular plant *Arabidopsis thaliana*. Specific Lhca and Lhcb paralogs are present in multiple copies in the *Physcomitrium patens* genome compared to one to two copies in *C. reinhardtii* and *A. thaliana*. The major antenna proteins encoded by Lhcbm also show greater redundancy and diversity in *Physcomitrium patens* (Iwai *et al*. 2018; Sun *et al*. 2023).

Comparing the genomes of *Physcomitrellopsis africana, F. hygrometrica* and *Physcomitrium patens* reveals both conservation and divergence of LHC genes. For example, *Physcomitrellopsis africana* has eight distinct Lhca genes, whereas *Physcomitrium patens* has 12 and *F. hygrometrica* has seven. Similarly, whereas *Physcomitrellopsis africana* has seven distinct Lhcb genes, *Physcomitrium patens* has 12, and *F. hygrometrica* has nine. Finally, *Physcomitrellopsis africana* has two distinct Lhcbm genes, *Physcomitrium patens* has 14, and *F. hygrometrica* has eight. However, echoing Kirbis *et al*. (2022), more light-harvesting complex (LHC) gene duplicates were retained in *Physcomitrium patens* than in *F. hygrometrica* and *Physcomitrellopsis africana* (Supplementary Fig. S3, Kirbis *et al*. 2022). Whereas *Physcomitrium patens* retained multiple LHC paralogs of possible WGD origin or gene duplications, *F. hygrometrica* and *Physcomitrellopsis africana* may have lost some of this gene redundancy. This pattern of differential retention is further supported by assessing the number of paralogs and orthologs for each LHC gene family across the three genomes (Supplementary Table S4). Although some copies of LHC genes were lost in *F. hygrometrica* and *Physcomitrellopsis africana*, key STN7 and STN8 kinases involved in photosynthetic acclimation are conserved and retained in all three genomes, suggesting retention of core light signaling components.

### Contaminant filtering and the identification of horizontal gene transfer events

In plants, horizontal transfer of genes, can occur at all phylogenetic depths, via introgression following hybridization between plant species (Pfennig 2021) to acquisition of genetic material from other domains (i.e., bacteria) or phyla of eukaryotes (e.g., fungi) (Soucy *et al*. 2015; Husnik and McCutcheon 2018). Identifying such transfers and hence the donors, is challenged by the occurrence of the plant’s microbiome (Koutsovoulos *et al*. 2016; Francois *et al*. 2020) and hence requires critical screening of sequencing outputs. Filtering before and after the assembly of *Physcomitrellopsis africana* was employed, illustrating that iterative approaches at all stages enhanced the final quality of the reference. Here, metagenomic tools (Centrifuge; Kim *et al*. 2016), optimized for long reads, identified contaminants before assembly. Since this initial filter did not include the short reads or assess fungal contributions in either sequence set, further filtering was conducted after assembly and annotation. An estimated 22% of the genes were of fungal origin and 31% were of bacterial origin. Scaffolds that contained numerous genes originating from algae, bacteria, or fungi (ABF) were removed. Separately, the annotated gene space was aligned to a set of “donor” and “recipient” databases. This identified 31 potential horizontal gene transfer (HGT) events. Following the set of best practices outlined by Francois *et al*. (2020), an additional set of 10 scaffolds was found to be contaminated as the flanking genes of the candidates were not of plant origin. This resulted in their removal (HGT filter). Two unique HGT candidates survived these filters, and the final scaffolding (Fig. 4).

**Fig. 4.**
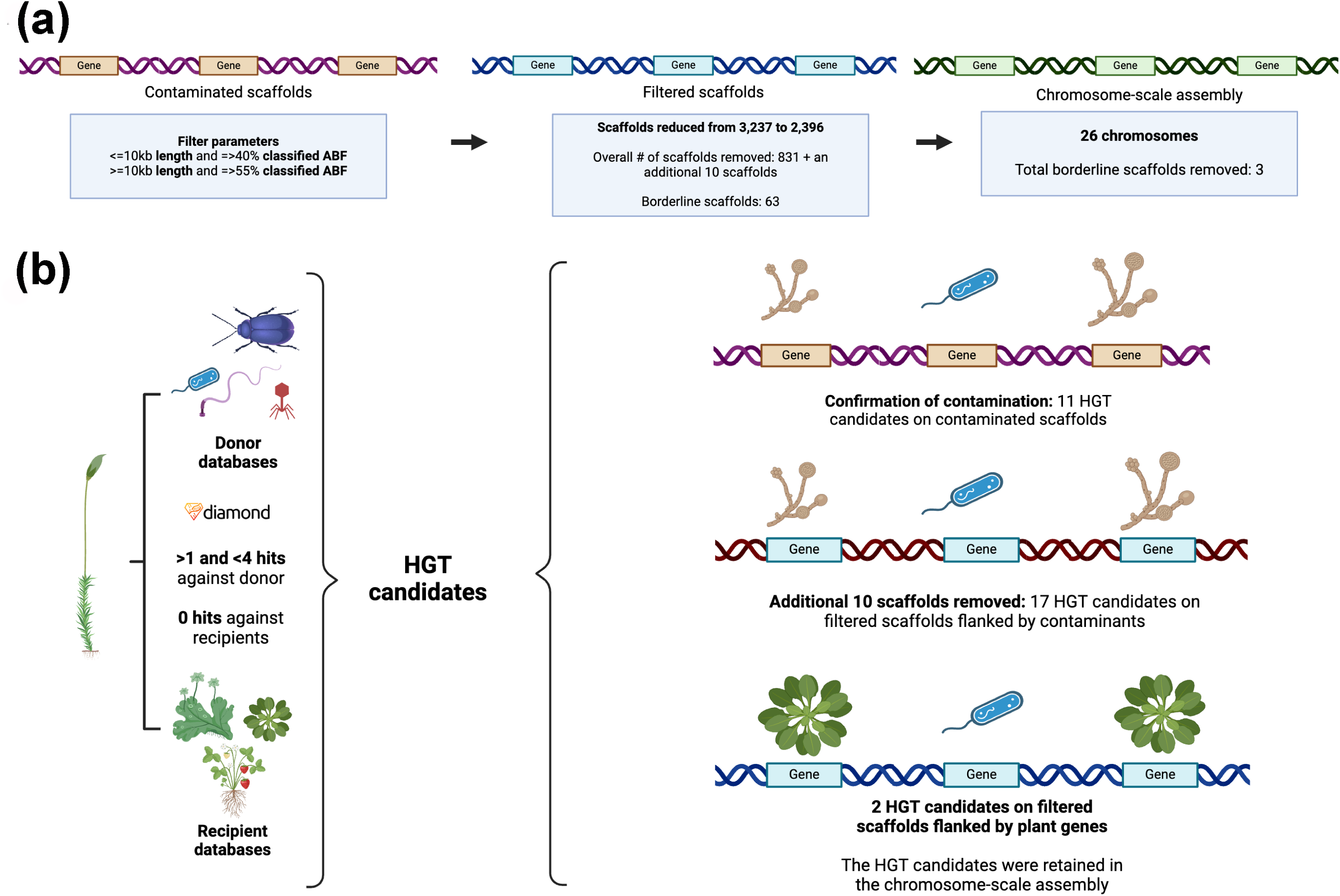
a) Contamination versus horizontal gene transfer (HGT) in the *Physcomitrellopsis africana* genome. Parameters used for removing a set of contaminated scaffolds from the draft genome based on functional characterization of the annotated gene space. Scaffolds with a length of 10 kb or less, and those with 40% or more of their total genes classified as archaea, bacteria, or fungi were removed. Additionally, scaffolds with a length greater than or equal to 10 Kb and having 55% or more genes classified as archaea, bacteria, or fungi were also excluded. In total, 831 scaffolds were removed because of this filtering process. Sixty-three scaffolds contained at least one gene annotated as a contaminant but did not meet the original thresholds for removing full scaffolds. When the chromosome-scale assembly was generated with RagTag scaffolding, three of these scaffolds were not placed in the 26 chromosomes. b) Identification of HGT candidates via sequence similarity comparisons of *Physcomitrellopsis africana* proteins against both “donor” databases (archaea, bacteria, fungal, and metazoan) and “recipient” databases (Streptophyta, Tracheophyta, and Spermatophyta). Proteins with >1 and <4 hits against all donor databases, and no hit against recipient databases were labeled as HGT candidates. This analysis was conducted on the contaminated scaffolds removed in A, confirming their contamination status. On the post-filtered scaffolds, there were some putative HGTs that were flanked by contaminants. These scaffolds were also removed from the assembly (additional 10), resulting in the retention of two HGT candidates in the final analysis.

The first HGT candidate, *Pa1_19801.1* (alignment to C-type lectin mannose-binding isoform-like XP_032077963.1). C-type lectins are the most frequent binding sites across all plants and animals. In vertebrates, CTLs have a function of pathogen recognition. HGTs in the context of plant defense and immunity are not new and have been identified before (Yue *et al*. 2012; Ma *et al*. 2020). In fact, Ma *et al*. (2020) identified a lectin that was acquired from bacteria. The second candidate, *Pa1_12964.1*, aligns to a hypothetical protein found in *Endogone* sp. FLAS-F59071 (RUS16308.1). While there is no information on the function of this gene, transfers from fungi are not surprising. As mentioned previously, fungal killer proteins KP4 have been acquired by mosses and play an important role in protonemal development (Guan *et al*. 2023).

These two candidates were manually validated by comparing alignments of de novo assembled transcriptomes. Both candidates are directly flanked by well-annotated plant (moss) genes. Additionally, aligning the full set of *Physcomitrium patens* and *Ceratodon purpureus* proteins to the *Physcomitrellopsis africana* scaffolds produced no alignments in proximity to the candidates, further suggesting that these HGT candidates are specific to *Physcomitrellopsis africana* (Fig. 5).

**Fig. 5.**
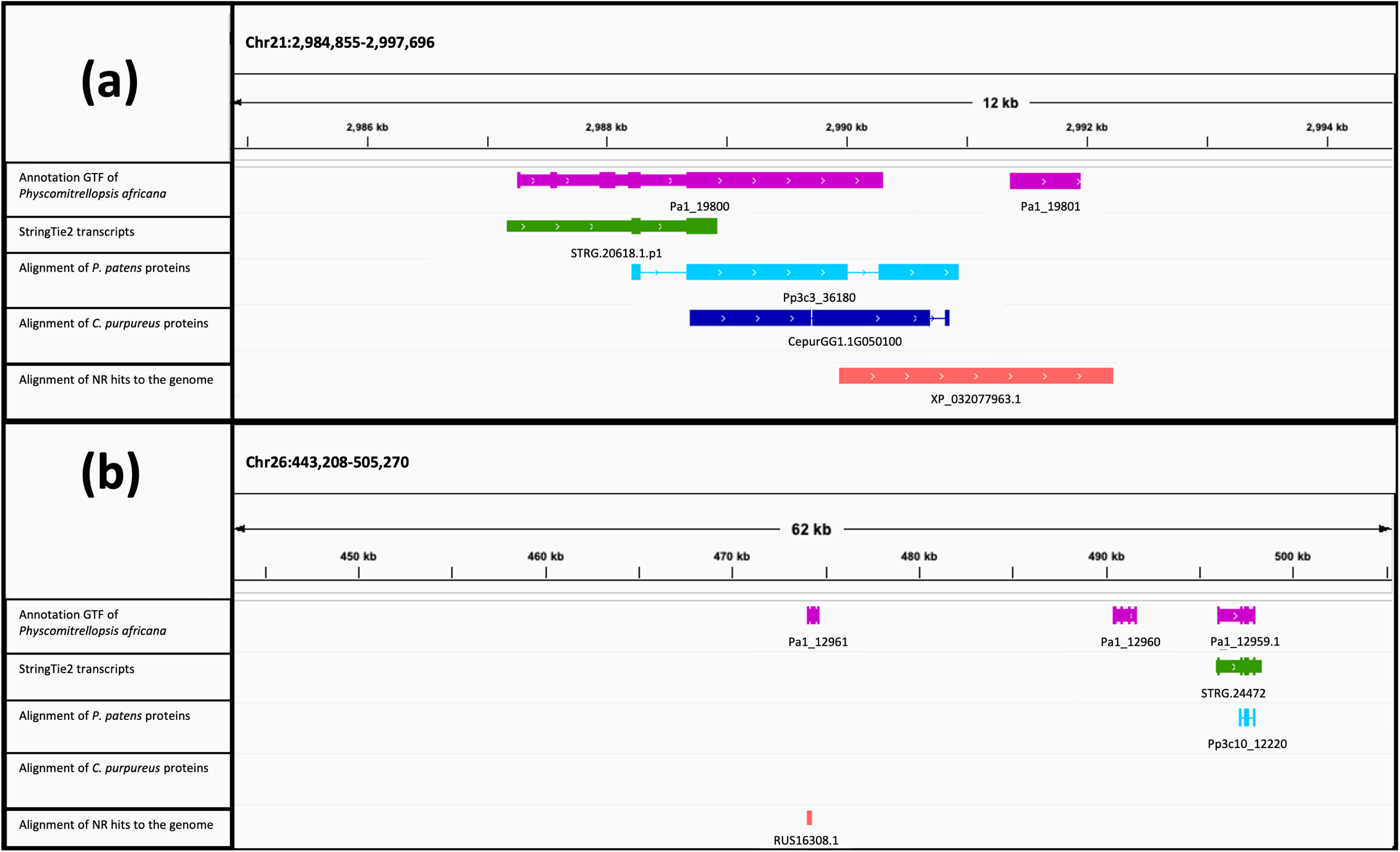
Integrated Genome Viewer (IGV) screens depicting five tracks of the *Physcomitrellopsis africana* genome. Track 1 shows the protein-coding structural annotation in context to the genome. Track 2 displays genome-guided transcript assemblies via StringTie2. Track 3 illustrates the alignment of the Horizontal Gene Transfer (HGT) candidate (nr database) to the genome. Tracks 4 and 5 show the alignments of *Physcomitrium patens* and *Ceratodon purpureus* proteins onto chromosomes 2 and 26. a) and b) show the HGT candidates *Pa1_04828.1* and *Pa1_22336.1*, where the HGT candidate alignment validates the presence of a protein. In both cases, independent RNA assemblies, via StringTie2, were not able to generate a supporting model. None of the other moss proteins align to the HGT candidates. c) highlights the example of *Pa1_15362.1*, where the StringTie2 transcript and the protein alignment reveal that the gene spans two gene models, indicating a false identification as an HGT candidate. Further transcriptomic evidence is required to modify the annotation model and establish the validity of these gene models.

#### Analyzing HGT candidates from Physcomitrium patens

The genomes of *Physcomitrellopsis africana* and *F. hygrometrica* were screened for the 273 putative horizontally transferred genes (HGTs) previously identified in *Physcomitrium patens* (Yue *et al*. 2012; Ma *et al*. 2022). Ninety-one of these genes (33%) were not found in the other two species, whereas 15 (5%) were shared only by *Physcomitrellopsis africana* and *Physcomitrium patens*, and seven (3%) only by *Physcomitrium patens* and *F. hygrometrica* (Supplementary Fig. S4). The greater number of shared HGTs between *Physcomitrium patens* and *Physcomitrellopsis africana* likely reflects their more recent divergence (at least 40 MYA) compared to that of *Physcomitrium patens* and *F. hygrometrica* (at least 60 MYA) (Medina *et al*. 2018; Bechteler *et al*. 2023). These findings suggest that horizontal gene transfer may have played a role in the evolution of mosses, potentially facilitating their adaptation to diverse environments over their long evolutionary history dating back 500 million years (Bechteler *et al*. 2023). The life cycle of mosses, with stages where cells are exposed to microbes, such as the egg or zygote within the open archegonium or motile sperm cells, could provide opportunities for the uptake of foreign genetic material from bacteria or fungi (Yue *et al*. 2012; Huang 2013). These vulnerable stages raise the hypothesis that putative microbial symbionts or associates of mosses may have been sources of horizontally transferred genes, potentially aiding mosses in adapting to harsh environmental conditions encountered as some of the earliest terrestrial plants.

## Data availability

All scripts and data are described in doi.org/10.5281/zenodo.11094703. NCBI BioProject ID PRJNA1020579 contains the genomic short reads and nanopore long reads (SRR26596311, SRR26596310 and SRR26666216) and the RNA reads (SRR26586950), as well as the whole genome assembly (GCA_036785485.1). The *de novo* assembled transcripts and annotation files are available at dx.doi.org/10.6084/m9.figshare.25724079.

## Acknowledgements

The authors would like to thank the Institute for Systems Genomics (ISG) at the University of Connecticut, including the Center for Genome Innovation for sequencing support and the Computational Biology Core for software support and access to high-performance computing.

## Funding

This study was made possible through the US National Foundation grants DEB-0919284 (fieldwork), DEB-1753811 to BG, and DBI-1943371 to JW.

## Conflict of interest

The authors declare no conflict of interest.

## Author contributions

J.W. & B.G. designed the study. B.G., N.W. & T.H. conducted fieldwork to sample the wild population. N. R., Y. C. & Y. L. generated genomic and transcriptomic data. A.S. & V.S.V. conducted all analyses. V.S.V., J.W. & B.G. wrote the paper. All authors approved of the final version.

## Notes

https://gitlab.com/PlantGenomicsLab/physcomitrellopsis-africana-genome

### Competing Interest Statement

The authors have declared no competing interest.

### Summary of Updates

The genome was brought up to chromosome-scale, All figures revised.

