## Supplementary material for "Crossroads of assembling a moss genome: navigating contaminants and horizontal gene transfer in the moss *Physcomitrellopsis africana*": Table1

**Table 1.** Genome annotation statistics for *Physcomitrellopsis africana*.

| **Annotation Method** | **Total genes** | **BUSCO (viridiplantae)** | **Mono: multi ratio** | **Annotation rate** |
| --- | --- | --- | --- | --- |
| BRAKER (RNA) | 60,917 | C:95.8%[D:15.3] | 0.52 | 65% |
| BRAKER (protein) | 37,752 | C:46.1%[D:16.0] | 0.95 | 73% |
| TSEBRA (BRAKER RNA + BRAKER protein) | 45,737 | C:91.3%[D:15.1] | 1.02 | 74% |
| BRAKER (RNA) + InterProScan filter + scaffold contam filter) | 23,561 | C:93.8%[D:13.6%] | 0.08 | 78% |
| BRAKER (RNA) + InterProScan filter + scaffold contam filter + HGT filter) | 23,535 | C:88.5%[D:11.3%] | 0.07 | 83% |
| Scaffolded to 26 chromosomes with RagTag | 22,925 | C:86.4%[D:11.1%] | 0.07 | 75% |
