## Supplementary for "Crossroads of assembling a moss genome: navigating contaminants and horizontal gene transfer in the moss *Physcomitrellopsis africana*"

**Supplementary Table S1.** Pre-QC Read Stats.

| Instrument | Total Reads | N50 (bp) | Average Read Length (bp) | Coverage (404 Mb) | Coverage (440 Mb) |
|---|---|---|---|---|---|
| Nanopore PromethION | 16,640,338 | 7,518 | 3,080 | 127 | 116 |
| Illumina Genomic Reads | 529,496,757 | N/A | 100 PE | 262 | 240 |

**Supplementary Table S2:** Post-QC Read Stats

| Read Type | Total Reads | N50 (bp) | Average Read Length (bp) | Coverage (404 Mb) | Coverage (440 Mb) |
|---|---|---|---|---|---|
| Nanopore PromethION | 14,495,188 | 4,561 | 2,233 | 80 | 73 |
| Illumina Genomic Reads | 359,619,533 | N/A | 100 PE | 178 | 163 |

**Supplementary Table S3.** Repeat content of the MaSuRCA genome assembly.

| | | | |
|---|---|---|---|
| **Sequences** | 3237 | | |
| **Total Length** | 502343981 bp | | |
| **GC Level** | 36.07% | | |
| **Bases Masked** | 253596074 bp | 50.48% | |
| | **Number of elements** | **Length (bp)** | **Percentage of sequence** |
| **Retroelements** | 182967 | 203774766 | 40.56 |
| **SINEs** | 116 | 50840 | 0.01 |
| **Penelope** | 44 | 43124 | 0.01 |
| **LINEs** | 5635 | 3528322 | 0.7 |
| **CRE/SLACS** | 25 | 87739 | 0.02 |
| **L2/CR1/Rex** | 0 | 0 | 0 |
| **R1/LOA/Jockey** | 0 | 0 | 0 |
| **R2/R4/NeSL** | 0 | 0 | 0 |
| **RTE/Bov-B** | 0 | 0 | 0 |
| **L1/CIN4** | 1520 | 1352049 | 0.27 |
| **LTR elements** | 177216 | 200195604 | 39.85 |
| **BEL/Pao** | 0 | 0 | 0 |
| **Ty1/Copia** | 7771 | 7313798 | 1.46 |
| **Gypsy/DIRS1** | 146546 | 183504589 | 36.53 |
| **Retroviral** | 0 | 0 | 0 |
| **DNA transposons** | 3075 | 2020920 | 0.4 |
| **hobo-Activator** | 161 | 85415 | 0.02 |
| **Tc1-IS630-Pogo** | 828 | 502079 | 0.1 |
| **En-Spm** | 0 | 0 | 0 |
| **MuDR-IS905** | 0 | 0 | 0 |
| **PiggyBac** | 0 | 0 | 0 |
| **Tourist/Harbinger** | 576 | 270022 | 0.05 |
| **Other** | 0 | 0 | 0 |
| **Rolling circles** | 7857 | 3140001 | 0.63 |
| **Unclassified** | 70182 | 35763560 | 7.12 |
| **Total interspersed repeats** | | 241559246 | 48.09 |
| **Small RNA** | 454 | 382891 | 0.08 |
| **Satellites** | 0 | 0 | 0 |
| **Simple repeats** | 177543 | 7162961 | 1.43 |
| **Low complexity** | 26138 | 1350975 | 0.27 |

**Supplementary Table S4.** Comparison of light-harvesting complex (LHC) gene complements in *Physcomitrellopsis africana*, *Funaria hygrometrica*, *Physcomitrium patens*, *Chlamydomonas reinhardtii**, and *Arabidopsis thaliana* (Alboresi et al., 2008).* The number of paralogs is shown for key LHC genes including antenna proteins (Lhca, Lhcb) and major light-harvesting proteins (Lhcbm). The asterisks (*) indicate that the numbers for *C. reinhardtii* and *A. thaliana* were taken directly from the published literature.

| **Gene** | ***Physcomitrellopsis africana*** | ***Funaria hygrometrica*** | ***Physcomitrium patens*** | ***Chlamydomonas reinhardtii **** | ***Arabidopsis thaliana ***** |
|---|---|---|---|---|---|
| Lhca1 | 3 | 2 | 3 | 1 | 1 |
| Lhca2 | 2 | 3 | 4 | 1 | 1 |
| Lhca3 | 2 | 1 | 4 | 1 | 1 |
| Lhca5 | 1 | 1 | 1 | 1 | 1 |
| Lhcbm | 2 | 8 | 14 | 9 | 0 |
| Lhcb3 | 1 | 1 | 1 | 0 | 3 |
| Lhcb4 | 1 | 2 | 2 | 1 | 4 |
| Lhcb5 | 2 | 1 | 2 | 1 | 5 |
| Lhcb6 | 1 | 2 | 2 | 0 | 2 |
| Lhcb7 | 1 | 1 | 1 | 1 | 1 |
| Lhcb9 | 1 | 2 | 2 | 9 | 2 |

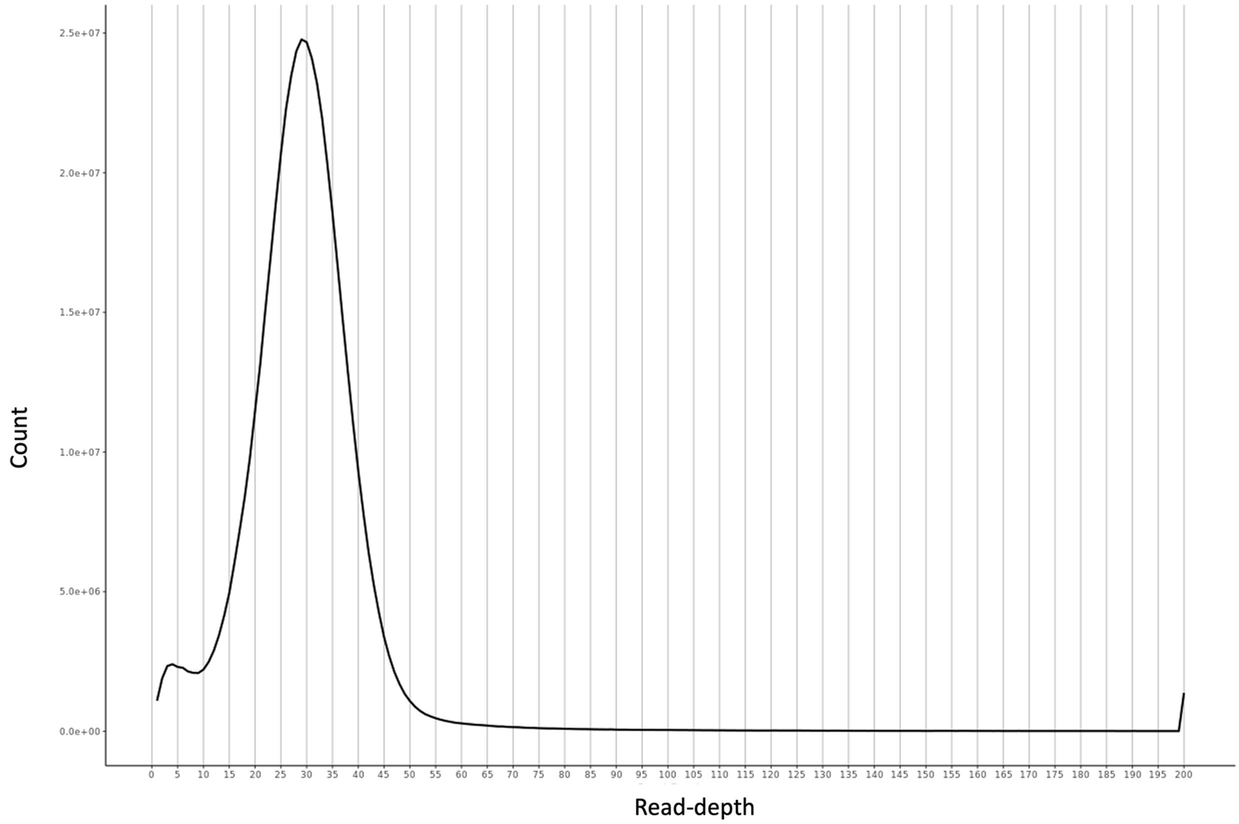

**Supplementary Fig. S1:** Histogram of k-mer coverage distribution for the *Physcomitrellopsis africana* genome assembly. The distribution shows two prominent peaks, with a smaller peak at ~7x coverage representing redundant haplotig sequences and a larger peak at ~65x representing the primary haploid genome coverage. The Purge Haplotigs pipeline used the local minima at 0x and 7x coverage as cutoffs to categorize and remove low coverage scaffolds presumed to be haplotigs or assembly artifacts. The 65x peak informed the coverage threshold for retaining the primary genome.

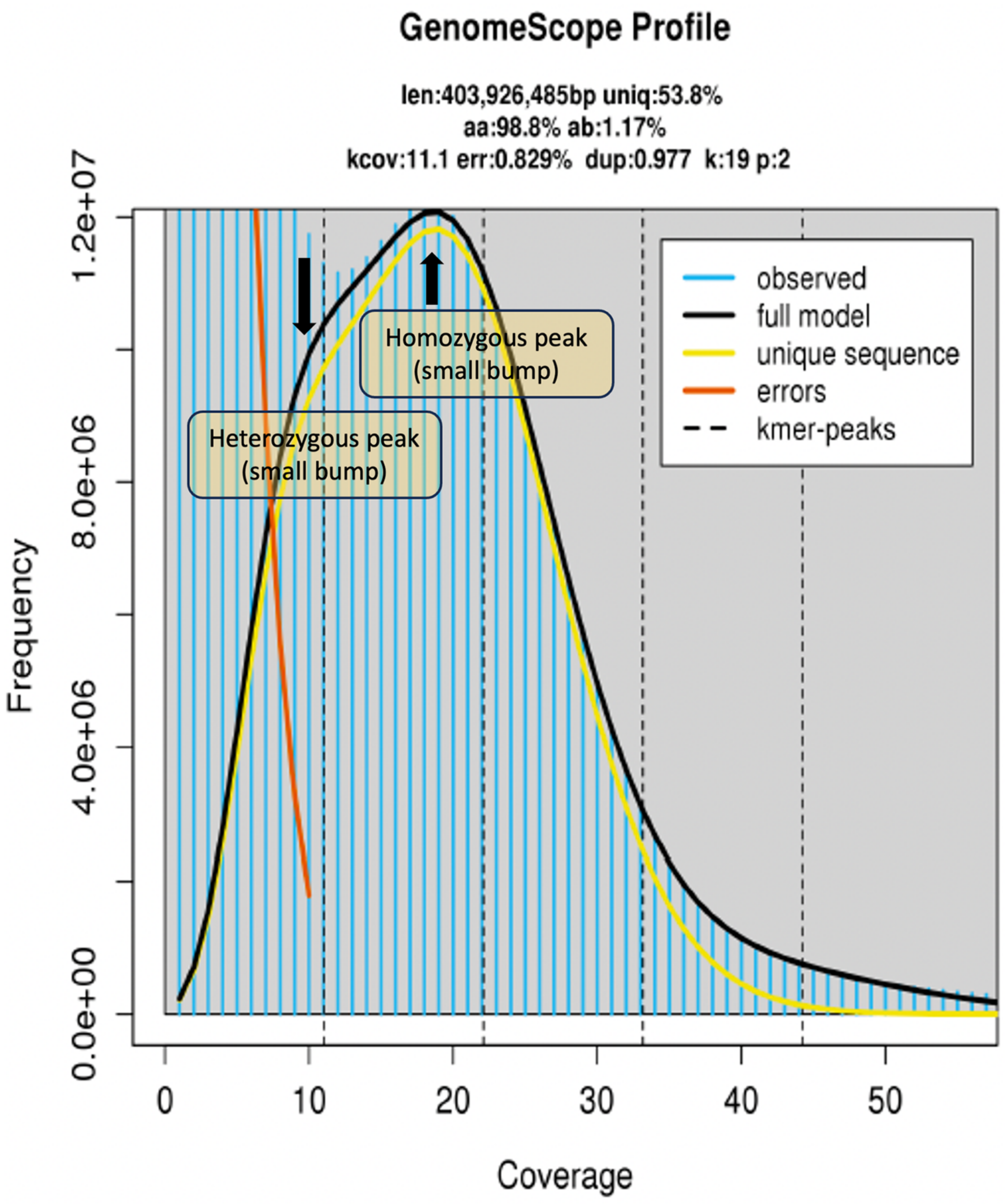

**Supplementary Fig. S2:** GenomeScope analysis of *Physccomitrellopsis africana* initially indicated an elevated heterozygosity rate (~1.17%) potentially reflecting the presence of sequence contaminants. Rigorous contaminant filtering procedures effectively mitigated this concern. Key genomic characteristics are summarized, including len (total genome length), uniq (unique genome percentage), kcov (kmer coverage), err (read error rate), and dup (read duplication rate). The peaks at 9X and 19X represent the estimated heterozygous and homozygous portions, respectively.

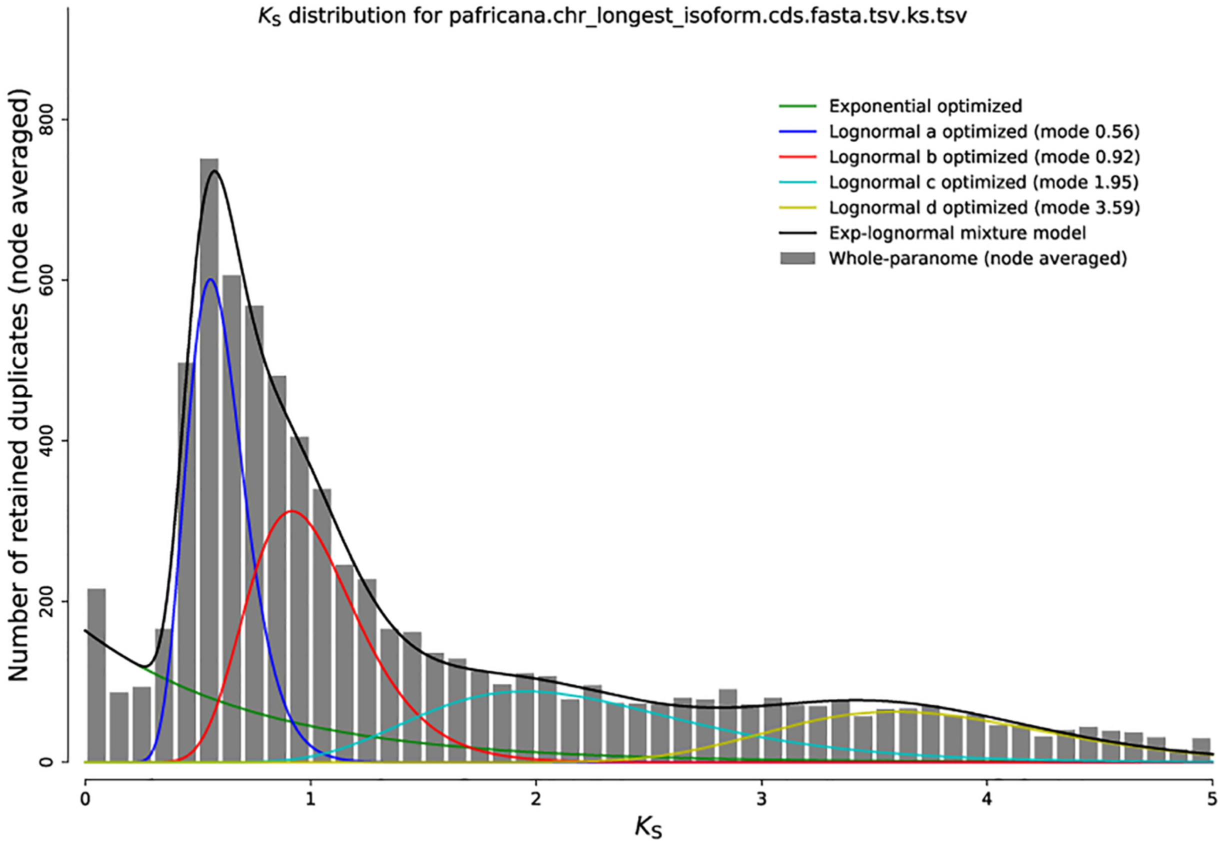

**Supplementary Fig. S3:** Ks density plots for *Physcomitrellopsis africana* reveal signatures of two ancestral whole genome duplications (WGDs), as evidenced by the two peaks. The x-axis shows Ks values representing synonymous substitutions per synonymous site. The y-axis shows the number of retained duplicates. The more recent WGD shows a peak at 0.56 (highlighted in blue), and the older WGD shows a peak at 0.92 (highlighted in red).

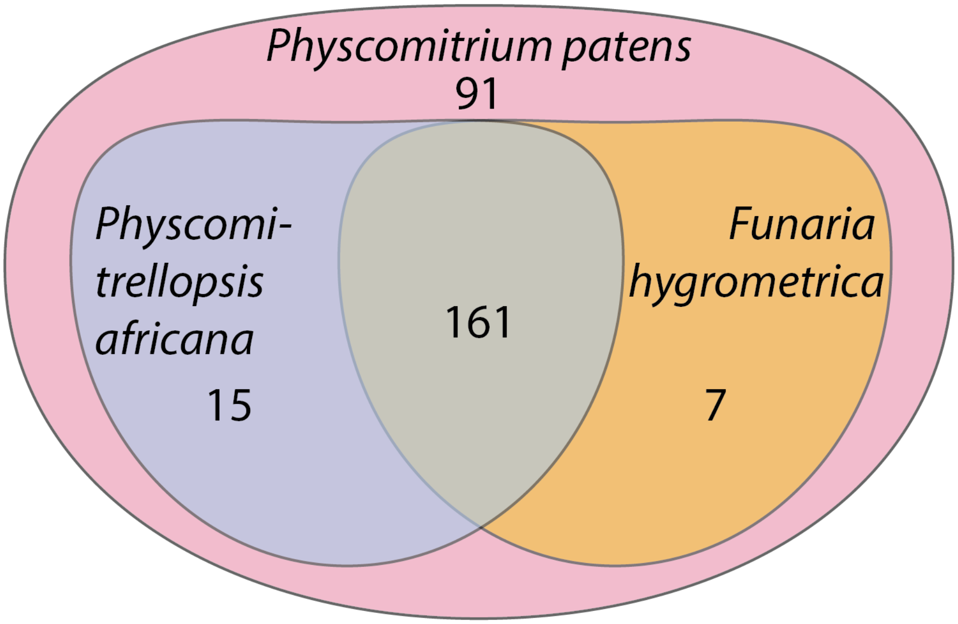

**Supplementary Fig. S4**. Distribution of the 273 horizontally transferred genes (HGTs) identified in *Physcomitrium patens* (Yue et al., 2012) in *Physcomitrellopsis africana* and *Funaria hygrometrica* genomes.
